# Geochemical records from loess sediments provide insights into early hominin influence on the landscape in the Khovaling region of Southern Tajikistan, Central Asia

**DOI:** 10.1101/2025.03.19.644098

**Authors:** Aljasil Chirakkal, Ekaterina Kulakova, Calin C. Steindal, Jago J. Birk, Anton Anoikin, Redzhep Kurbanov, Petr Sosin, Peixian Shu, David K. Wright

## Abstract

This study presents a multi-proxy investigation of loess–paleosol sequences from the Khovaling Loess Plateau (KLP) in southern Tajikistan to explore long-term human-environment interactions during the Early Paleolithic. This region has been the focus of numerous archaeological projects since Soviet times, as it contains evidence of some of the oldest (∼0.9 Ma) hominin occupations in Central Asia and is one of the key regions connecting eastern Asia to centers of hominin evolution that lie to the west. Previous archaeological expeditions have discovered rich collections of stone tools from alternating non-pedogenically modified loess and paleosol sequences at the archaeological sites of Karatau, Lakhuti, Obi-Mazar, and Kuldura. We analyzed lipid biomarkers such as *n*-alkanes, polycyclic aromatic hydrocarbons (PAHs), together with newly generated stable isotopes (δ¹³C, δ¹⁵N), magnetic susceptibility, and archaeological assemblages from three sites: Obi-Mazar/Lakhuti (on-site), and non-artifact-bearing strata from Kuldara (near-site) and Khonako-II (off-site), focusing on pedocomplexes (PCs) 4, 5, and 6 and their intercalated loess layers. High odd-over-even predominance (OEP) and average chain lengths (ACL) suggest that soil organic matter is predominantly derived from higher-order terrestrial plants, with better preservation in paleosols compared to loess layers. The inverse correlations of δ¹³C with TOC and C/N reflect organic matter degradation, while elevated δ¹⁵N in paleosols suggests enhanced nitrogen cycling under warmer and drier conditions. Magnetic susceptibility and δ¹³C trends reveal a progressive shift toward open grassland ecosystems since ∼0.8 Ma, with intensified pedogenesis during interglacial stages MIS 11, 13, and 15. The spatial distribution of PAH concentrations and lithic artifact densities further highlights human-environment interactions in the KLP locality. Obi-Mazar exhibits abundant lithic materials and high PAH levels, indicating repeated occupation and frequent burning in a resource-rich riverine setting. Kuldara exhibits moderate PAHs but minimal artifacts, whereas Khonako-II records only a minimal level of signals, reflecting a decrease in human impact with distance from the water source. Together, these findings demonstrate that Early Paleolithic hominins preferentially occupied ecologically stable, pedogenically developed zones during interglacial phases, contributing to localized fire regimes and shaping their landscapes. This study provides a refined paleoenvironmental framework for understanding hominin adaptations in Central Asia during the Pleistocene.

## 1. Introduction

Changes in past climate conditions that enabled the spread of grasslands in the ancestral homelands of hominins are interpreted as having been critical landscape-level circumstances associated with significant milestones in human evolution and adaptations, such as bipedal anatomical adaptations, technological innovation, and diversification of human populations (Cerling et al., 2011; Lupien et al., 2020). Archaeological/paleoanthropological studies and theoretical models have been developed to quantify these climate-human evolutionary relationships (Grove, 2014; Kingston, 2007; Lupien et al., 2020; Potts and Faith, 2015). Based on climate-human evolution models, Asia is subdivided into two main pan-archaeological regions: Westerlies Asia (i.e., the arid central region of SW China and Central Asia) and Monsoonal Asia (i.e., the eastern and southern regions). The behavior of humans, including long-term technological variability, occupation intensity, and livelihood activities during the mid-late Pleistocene, tends to bifurcate according to these macroregions (An et al., 2012; Caves et al., 2015; Chen et al., 2019; Dennell, 2013; Dodonov, 1991; Gao et al., 2017).

In turn, Central Asia is further divided into two regions that differ both climatically and in terms of the specifics of their archaeological industries. In the Caspian zone and the northern part of the region (Kyrgyzstan, Kazakhstan), assemblages are found that include hand axes and are considered by researchers within the framework of the Acheulean tradition (Davis and Ranov, 1999; Finestone et al., 2022; Glantz, 2011). However, in the southern part of the region (Tajikistan), Early Paleolithic sites belong to another cultural tradition, which lacks bifaces but contains a significant number of pebble choppers and unifacial tools (Fig. 1) (Ranov, 1995; Ranov et al., 1995; Ranov and Davis, 1979; Schäfer et al., 1998). The existing interpretations of hominin behavior from this region hypothesize that interglacial times were characterized by warm and wet periods with high precipitation, which resulted in intensified pedogenesis, resource-rich zones, and forest-grassland mosaics ripe for human exploitation across vast portions of Southern Tajikistan (Dodonov and Baiguzina, 1995; Mestdagh et al., 1999; Ranov, 1995; Yang et al., 2006). There are also several significant riparian zones with high-quality raw materials for knapping that are not common in the loess-covered landscapes of the region. These conditions supported the manufacture of expedient Paleolithic technologies, which is evidenced by the presence of several lithic-bearing archaeological sites within river valleys (Ranov and Davis, 1979; Schäfer et al., 1998). In contrast, glacial periods were characterized by greater aridity, cooler temperatures, and the spread of grassier environments (Pakhomov, 2006; Parviz et al., 2020), which had a less favorable environment for hominins based on the almost complete absence of lithic artifacts in the loess layers corresponding to glacial periods.

**Fig. 1.**
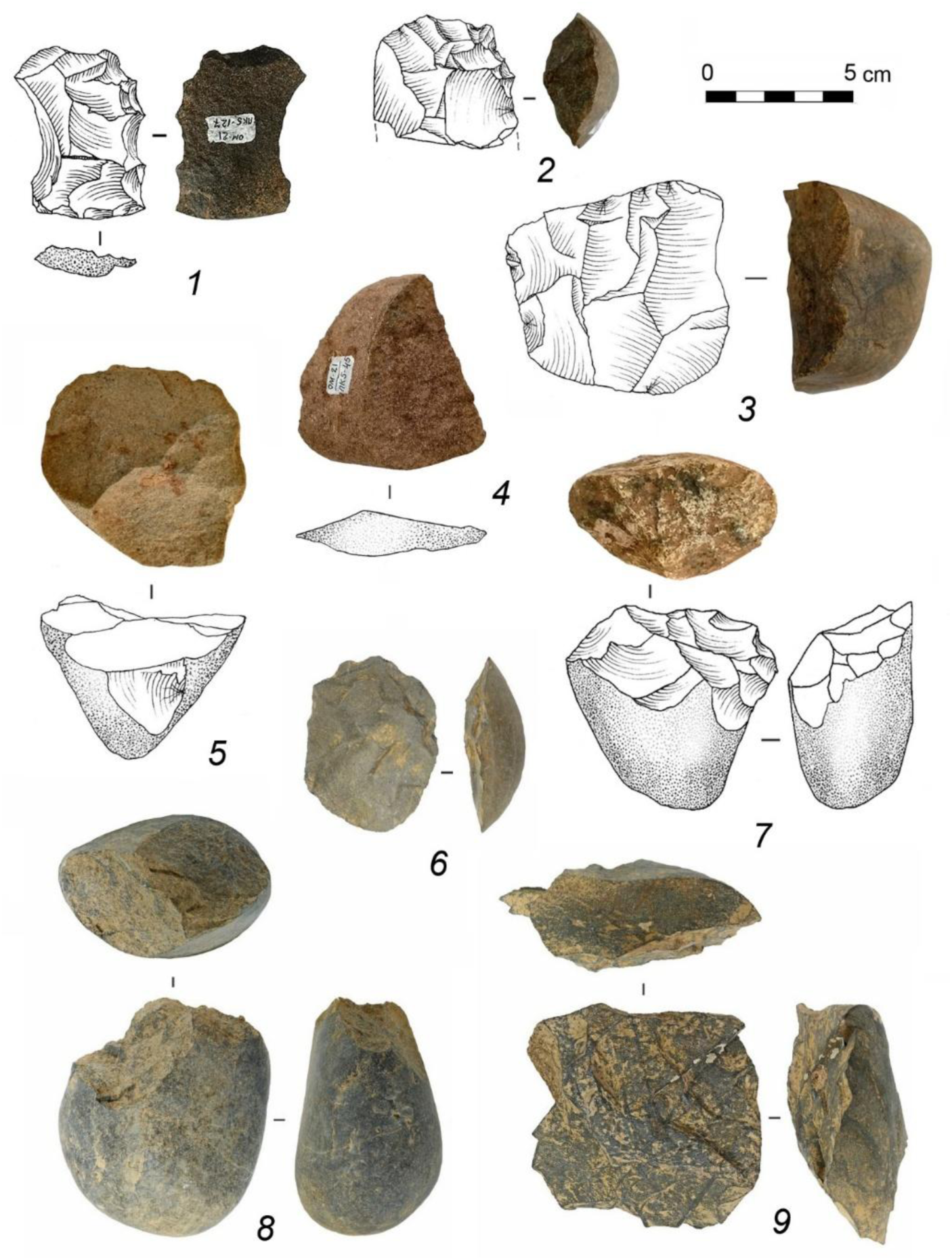
Types of lithic artifacts excavated from the KLP locality: 1, 6 -unifaces; 2 -denticulate tool; 3, 9 -radial cores; 4 -slice/”citrone”; 5 – slice core; 7, 8 -choppers.

The most representative Early Paleolithic technocomplex in Central Asia today is the Karatau culture, which existed in Tajikistan for approximately 200 ka (∼600 to 400 ka). The main archaeological assemblages from this period have been documented at the Obi-Mazar/Lakhuti sites in the Khovaling loess region (Fig. 2), with more than 4000 lithic artifacts recorded. The toolkits include choppers, pebble side-scrapers, simple side-scrapers, denticulate and notched tools, as well as unifaces (Fig. 1). The general analysis of the lithic assemblages associated with deposits from ∼600-400 ka is defined by the undustrial unity of both their preferred primary knapping techniques and composition of the toolkits. This technological stability, along with the association of assemblages with deposits of a single context (paleosols), suggests a potential link between the composition of the lithic industry and the environmental conditions in which the ancient hominins who produced these tools lived.

**Fig. 2.**
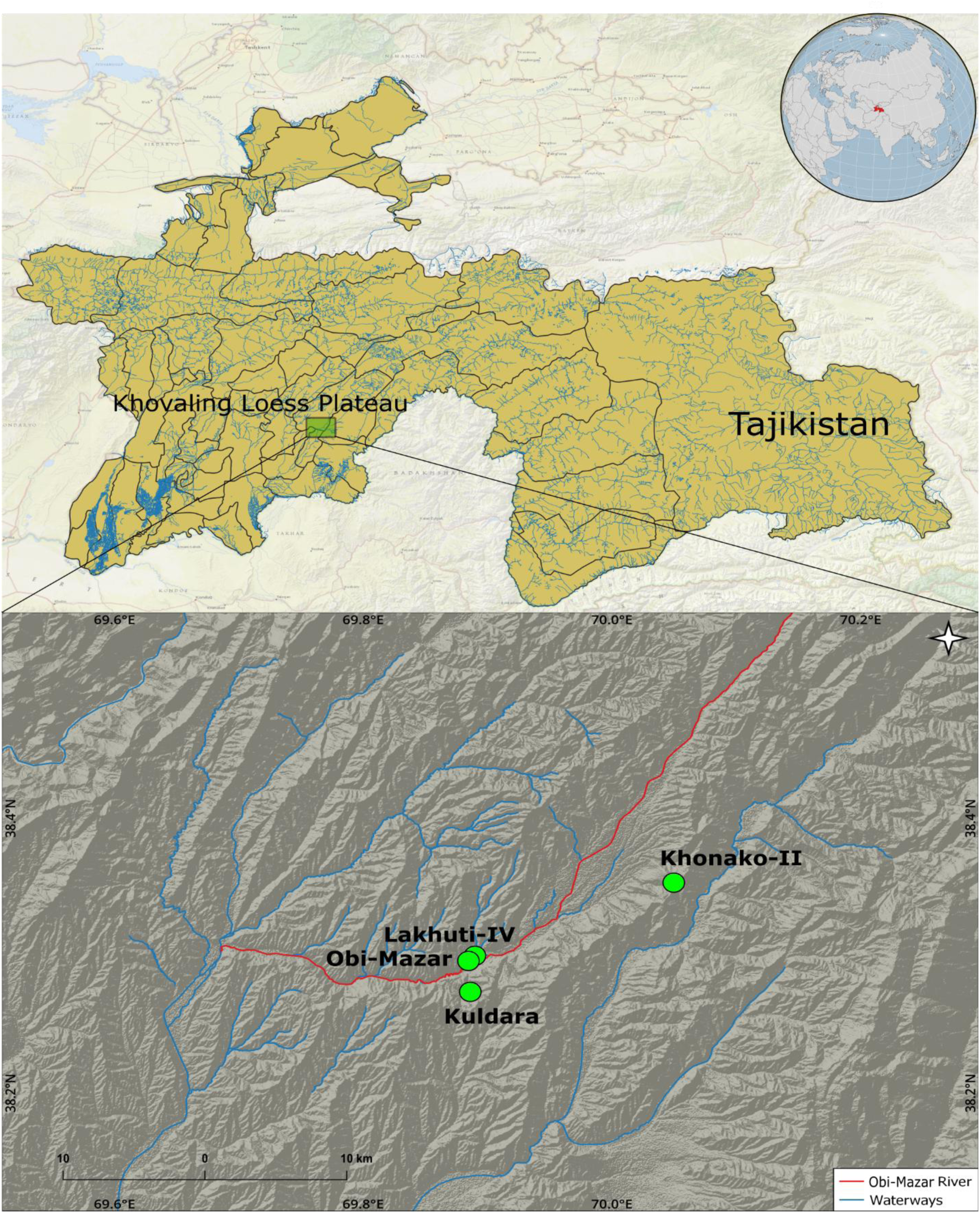
Location of Obi-Mazar, Lakhuti-IV, Kuldara, and Khonako-II sites on the Khovaling Loess Plateau in Southern Tajikistan, Central Asia.

Recent advances in the application of molecular biomarkers and biogeochemical indicators in archaeology have shaped our understanding of key aspects of past ecological and climatic dynamics associated with humans (Brittingham et al., 2019; D’Anjou et al., 2012; Harrault et al., 2022; Schatz et al., 2011; Stancampiano et al., 2023). Lipid biomarkers, such as leaf wax-derived alkanes (*n*-alkanes) and pyrogenic polycyclic aromatic hydrocarbons (PAHs) are well preserved in sedimentary organic matter (SOM) for hundreds to thousands of years and have been frequently used in paleoenvironmental studies (Ankit et al., 2022; Badewien et al., 2015; Topness et al., 2023; Vachula et al., 2019, and references therein). The long chains of *n*-alkanes (C_27_-C_33_) give insights about past vegetation changes (Zhang et al., 2006), while pyrogenic PAHs (e.g., pyrene) provide direct evidence for the past burning of vegetation in the catchment (Lima et al., 2005). In addition to these molecular markers, stable carbon (δ^13^C) and nitrogen (δ^15^N) isotopic records in SOM provide valuable palaeovegetational evidence regarding C_3_/C_4_ vegetation (Hatté et al., 1998; Liu et al., 2005) and climatic conditions associated with land use changes (Eshetu and Högberg, 2000).

While previous studies from the same sequences focused on lipid biomarker signatures (Chirakkal et al., 2025a,b), the present study introduces new datasets on stable isotopes (δ¹³C, δ¹⁵N), TOC, TN, and magnetic susceptibility. These geochemical proxies are integrated with the previously published lipid biomarker data to reconstruct paleoclimate, vegetation changes, and their relationship with early human occupation from three loess-paleosol sequences at the Khovaling loess region in Southern Tajikistan (Central Asia). A combination of these novel palaeoecological proxies has previously been applied to loess-palaeosol sequences (e.g., Schatz et al., 2011; Tarasov et al., 2013; Zech et al., 2012) to reconstruct Quaternary palaeoenvironments. The present study is the first one to use these novel palaeoecological proxies to investigate the potential early human impacts on the environment at archaeological sites in Khovaling, Tajikistan.

## 2. Study area

The Khovaling Loess Plateau (KLP) of Southern Tajikistan is a promising area to contribute to a broad understanding of Eurasian human evolution and migration through the reconstruction of vegetation regimes associated with Early-Middle Pleistocene *Homo* species occupations. From an ecological and climatological perspective, the KLP consists of loess-paleosol sequences with sufficient thickness for a thorough study of paleoecological changes that occurred during the accumulation of wind-blown loess deposits, and its mid-continental location reflects linkages with global climate events (Buylaert et al., 2024; Challier et al., 2024; Meshcheryakova et al., 2023). These loess-paleosol sequences consist of alternating thick non-pedogenically modified loess (L) units, resulting from high dust activity during glacial periods, and paleosols or pedocomplexes (PCs) developed during interglacial periods under warmer and wetter climatic conditions. Our study locality is defined by four loess-paleosol sequences situated around the valley of the Obi-Mazar River in the Khovaling district of Southern Tajikistan (Fig. 2). In this context, archaeological layers are easily identified because the sedimentary matrix consists exclusively of silt-or clay-sized particles (Tokareva et al., 2025) and there are no proximal colluvial or alluvial sources of stones, meaning that large clasts are, by definition, manuports or artifacts. Therefore, the distribution and concentration of lithic artifacts serve as indicators of human activity and occupation intensity during the time intervals recorded in the sequences (Anoikin et al., 2023; Ranov, 1995; Schäfer et al., 1998).

The first study location is Obi-Mazar (OBM), which is positioned in the middle of the Obi-Mazar/Lakhuti outcrop (38.28° N, 69.88° E) (Figs 2 and 3a), and also includes seven pedocomplexes, which have a thickness of 90 m from the bottom of PC 7 to the Holocene soil. At the end of the 20th century, excavations at the OBM-IV and -VI sites uncovered a substantial collection of lithic artifacts (about 1,500 items; Table 1). However, significant portions of OBM were later destroyed by a massive landslide. The modern OBM section is more than 100 m inland from the original sites and represents their periphery (Anoikin et al., 2023). The second location is Kuldara (KUL), which is situated in Kuldara gully ∼2 km away from the Obi-Mazar/Lakhuti outcrop and includes up to 12 pedocomplexes and has a loess depth of 85 m (38.27° N, 69.88° E) (Fig. 2 and 3b). It is worth noting that Kuldara gully is a minor tributary of the Obi-Mazar River, stretching for 5-6 km. This may explain the relatively small collection of lithic artifacts and the apparent low intensity of occupation in antiquity. A few stone tools (Table 1) were discovered from PCs 4, 5, 6, while this site is more known for the oldest lithic artifact findings (40 items) in the region in PCs 11-12 (Ranov, 1995). The third study location is Khonako-II (KH-II), situated on the Khonako outcrop, which is ∼15 km north-east of the group of other sites (38.35 ° N, 70.04° E) (Fig. 2 and 3c). KH-II is a geological sequence with 180 m thickness and consists of up to 24 pedocomplexes (Dodonov, 1986). The KH-II section contains almost no archaeological material, with only a single artifact found in PC 4 (Table 1). It should be noted that the site of KH-III, located 100 m from KH-II, 25 artifacts were recovered from PC4 by Schäfer et al. (1998). However, the interpretation of these finds was that they were highly dispersed (Schäfer et al. 1998) and likely represent the remains of ephemeral hunting camps or lookout areas, more so than settlements. Given this, and with the low amount of archaeological material (n=1) from KH-III PCs 4-6 recovered from the 2022 and 2023 campaigns, we classify these units as archaeologically sterile.

**Fig. 3.**
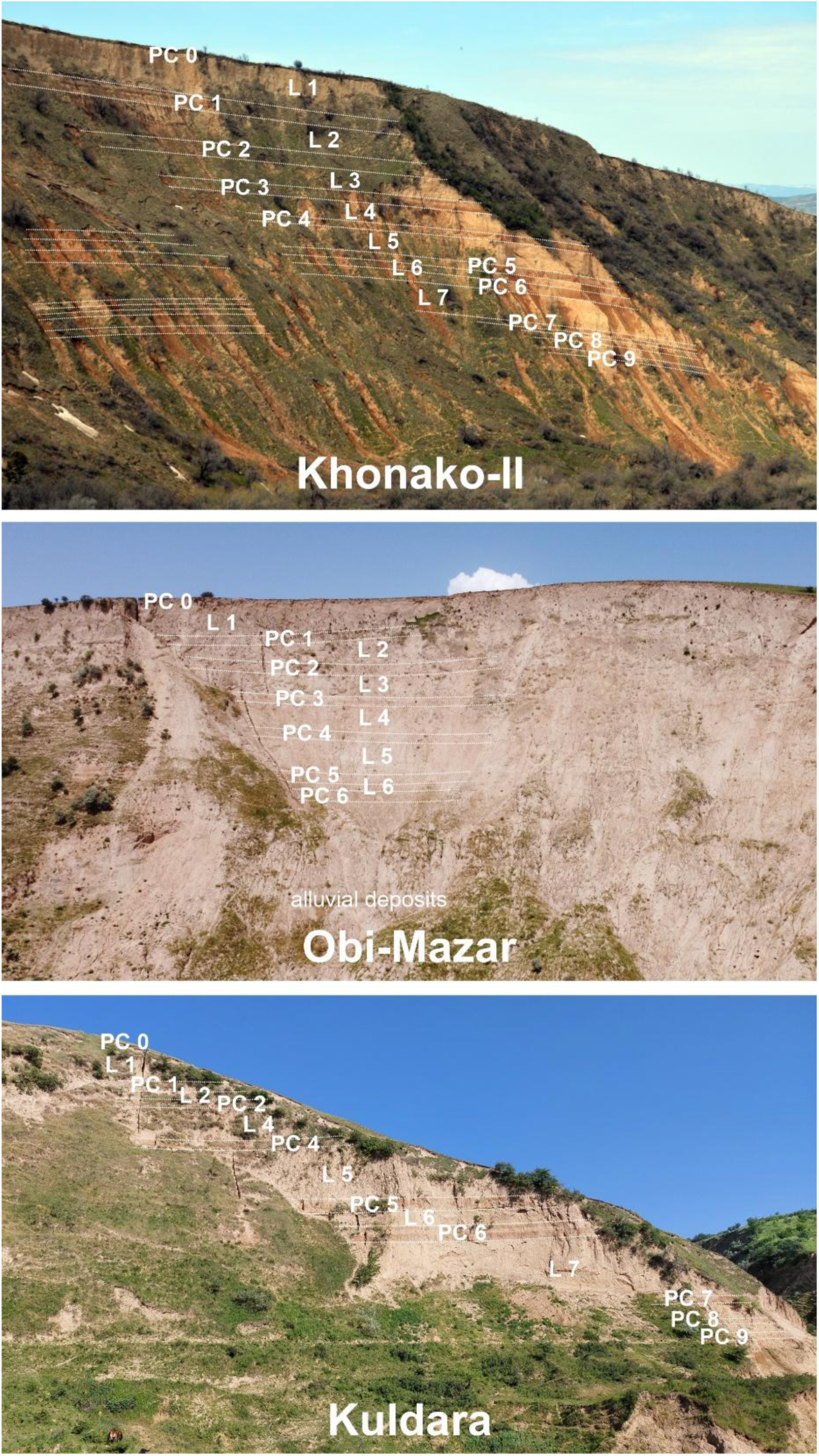
Field photograph of stratigraphy and general view of the (a) Obi-Mazar (OBM), (b) Kuldara (KUL), and (c) Khonako-II (KH-II). PC: pedocomplex, L: Loess.

**Table 1.** The total number of lithic artifacts found at KLP locations, according to the previous findings by Anoikin et al. (2023), Ranov (1995), and Schäfer et al. (1998).

| Study location | Obi-Mazar (on-site) | Kuldara (near-site) | Khonako-II (off-site) |
| --- | --- | --- | --- |
| PC 4 | 2125 | 2 | 1 |
| PC 5 | 247 | 1 | 0 |
| PC 6 | 331 | 1 | 0 |

Modern climate data indicate that study locations in the Khovaling region of Southern Tajikistan experience a Mediterranean-type climate, characterized by dry summers and precipitation primarily occurring from early winter to spring. The modern mean annual temperature in Khovaling is 12-13°C, but significant seasonal temperature fluctuations occur, with variations exceeding 20°C between May and September due to its continental location (Fig. S1a). Most of the precipitation (∼663 mm) falls between October and June, with less than 30 mm during June to September (Fig. S1b). The precipitation in the region is driven by Mediterranean cyclones and the Siberian Winter High, with moisture coming either directly from the Mediterranean or the North Atlantic Ocean, or recirculating from the Black and Caspian seas (Aizen et al., 2006; Li et al., 2019). The Pleistocene climate of Tajikistan was shaped by a combination of long-term aridification and the cyclic oscillation between glacial and interglacial periods, driven by global climatic shifts and regional weather systems. The uplift of the Tibetan Plateau and associated orogenesis in the Tien Shan, Pamir, and Hindu-Kush mountain ranges played a crucial role in restricting moisture penetration into Tajikistan from monsoonal systems (Li et al., 2019; Miao et al., 2012). During glacial periods, intensified aridity, reduced vegetation cover, and stronger eolian activity contributed to extensive loess deposition, with continental high-pressure systems limiting the intrusion of westerlies (Ding et al., 2002). In contrast, interglacial periods have a greater influence from North Atlantic-Mediterranean westerlies, resulting in more humid conditions (Dodonov and Baiguzina, 1995).

## 3. Materials and methods

### 3.1 Sampling strategy

During the 2021 and 2022 fieldwork seasons, OBM, KUL, and KH-II loess-paleosol sequences in KLP were targeted for pedostratigraphic analysis and sampling. The mapping and sampling strategies employed were identical for these study locations and consisted of procedures outlined in Chirakkal et al. (2025a) following standard geoarchaeological techniques using the United States Department of Agriculture methods (Schoenberger et al., 2012). The sampling of each study site was performed by sampling trenches that were cut into exposed profiles. The trenches were cut back from the modern surface by >50 cm to avoid contamination by modern organic matter and surface runoff. The lipid biomarkers and stable isotope samples were collected from PCs 4, 5, and 6 from three study locations. Here, we defined a “site” as being a distinct location within the loess-paleosol sequences that contains evidence of past human presence, specifically through the discovery of lithic artifacts. Based on this, we separated these study locations into “on-site” (OBM), “near-site” (KUL), and “off-site” (KH-II) contexts based on the recovery of lithic artifacts (Table 1) and distance from the Obi-Mazar River. The sediment samples of ∼50 g for lipid biomarker and stable isotope analysis per sampling point were retrieved in high-density polyethylene Whirl-paks using a trowel or knife, avoiding contact with human hands. All sample bags were kept open for drying at room temperature for >24 hours before repacking. Sample bags were checked daily to ensure that humidity did not accumulate and were further dried in a laboratory convection oven at 40 °C for >72 hours.

### 3.2 Magnetic susceptibility

Field magnetic susceptibility measurements were performed for the KUL and OBM localities using a portable PIMV kappameter (Geodevice, Russia) with a sensitivity of up to 10^-7^ SI units. The measurement was carried out continuously over the entire thickness of sampling sections at the locations, the average step was about 3.5 cm. Measurement of the field magnetic susceptibility for the KH-II locality was incomplete, so for this location, we provide the data of mass-specific magnetic susceptibility obtained from bulk samples. These loose, unoriented samples were collected at a 2-cm interval every 10 cm for paleosols and 20 cm for loess layers and air-dried in a laboratory for further measurements. Mass normalized magnetic susceptibility was measured using MFK1-FA kappabridge (AGICO, Czech Republic) at a frequency of 976 Hz and a field intensity of 200 A/m. The comparison of the field magnetic susceptibility and mass-specific measurements for the same locations (for eg, KH-II and KUL) showed identical behavior.

### 3.3 Stable isotope analysis

All samples for the study of δ^13^C and δ^15^N isotopes were homogenized using a sterile agate mortar and pestle. The sediment samples destined for analysis of δ^13^C were saturated in 1M HCl for >24 hours to remove carbonates, and the step was repeated if bubbles were still present. When the reaction with acid was stopped, distilled water was poured, which was added 3 times to the tubes, agitated, and centrifuged to complete the removal of carbonates and acid. Then samples were kept in an oven at 40°C with caps open until they were thoroughly dried. The samples for δ^15^N measurements were not acidified. The δ^13^C and δ^15^N samples were weighed and packed into sterile tin capsules. The elemental concentrations of C and N, as well as δ^13^C and δ^15^N values, were analyzed using a Flash 1110 Elemental Analyser (EA) with a no-blank autosampler connected to a Delta V Plus isotope ratio mass spectrometer (Thermo Fisher Scientific). The standards utilized were IAEA-N-1 and IAEA-N-2 for N isotopes and USGS-24, IAEA-600, and IAEA-CH-6 for C isotopes (https://analytical-reference-materials.iaea.org/). The precision of standards was <0.2 per mil ‰ for both δ^13^C and δ^15^N. The ^13^C and ^15^N composition of a sample relative to an international standard (VPDB, atmospheric N_2_) is expressed in ‰ in the international delta (δ) notation according to the following equation:

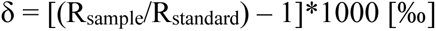

where R is the ratio of the heavier to the lighter isotope of carbon and nitrogen (^13^C/^12^C and ^15^N/^14^N).

### 3.4 Lipid biomarker analysis

The *n*-alkane and PAH data from the loess–paleosol sequences of OBM, KUL, and KH-II have been published previously (Chirakkal et al., 2025a, 2025b). These datasets are re-examined here in conjunction with new stable isotope (δ¹³C, δ¹⁵N), total organic carbon (TOC), total nitrogen (TN), and magnetic susceptibility data to develop an integrated interpretation of paleovegetation dynamics, pedogenesis, and human-environment interaction. The total lipid extraction and separation procedures have been previously reported in Chirakkal et al. (2025a, 2025b). In brief, approximately 15 g of dried, crushed sediment samples were extracted using a Soxhlet extractor with a solvent mixture of dichloromethane (DCM) and methanol (MeOH) (2:1, v/v). The total lipid extracts were concentrated and further separated into non-polar and polar fractions by silica gel column chromatography. The non-polar fractions were eluted with a solvent mix of hexane and DCM (4:1, v/v), and polar fractions were eluted with DCM and MeOH mixture (1:1, v/v), respectively. The fractionated non-polar aliquots were analyzed by gas chromatography-mass spectrometry (GC-MS, Shimadzu). The samples were injected by an autosampler in pulsed splitless mode. The GC is equipped with a DB-5MS capillary column (30 m × 0.25 mm i.d. × 0.25 μm film thickness). Helium was used as the carrier gas at an average velocity of 40 cm/sec. The ion source operated in the electron ionization mode at 70 eV. The initial temperature of the GC oven was programmed to 60°C and then ramped to 320°C (isothermal for 10 min) at 3°C/min. Total ion chromatogram (TIC) and selected ion monitoring (SIM) were produced and desired compounds were identified by comparing the elution pattern and mass spectra from compound peaks of the external standards and the NIST library.

The *n*-alkane based proxy OEP, along with ACL and the C_27_+C_29_/C_31_+C_33_ ratio derived from previously published datasets (Chirakkal et al., 2025a, 2025b) are incorporated for palaeoenvironmental reconstructions in conjunction with newly generated stable isotope and magnetic susceptibility data.

Odd − over − even predominance (OEP) = 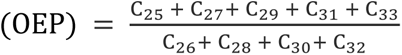 (Scalan and Smith, 1970)

## 4. Results

### 4.1 Age model and stratigraphic correlations

The chronostratigraphic correlation of Tajik loess to the Chinese loess and deep-sea oxygen records has shown that the first nine pedocomplexes in Southern Tajikistan are within the Brunhes chron, where PC 1 corresponds to Marine Isotope Stage (MIS) 5, each subsequent pedocomplex (PC 2-7) corresponds to the subsequent interglacial. The weakly developed PC 8 formed during a warm substage of the glacial stage MIS 18 (Fig. 4). The first paleomagnetic data found the Brunhes-Matuyama boundary in the glacial loess between PC 9 and 10 (Penkov and Gamov, 1980). However, recent detailed studies confidently place it at the base of PC 9 (Kulakova and Kurbanov, 2023), serving as a reliable time control point for the correlation of PC 9 with MIS 19. The current age model of Tajik loess was verified by recent geochronological research of the uppermost PCs using Optically Stimulated Luminescence (OSL) to confirm the formation of PC 1 and PC 2 during MIS 5 (71-130 ka) and MIS 7 (191-243 ka), respectively (Buylaert et al., 2024; Challier et al., 2024). Based on this age model, PC 4 corresponds to MIS 11 (364 – 427 ka), PC 5 to MIS 13 (474 – 528 ka), and PC 6 to MIS 15 (563 – 621 ka) (Fig. 4).

**Fig. 4.**
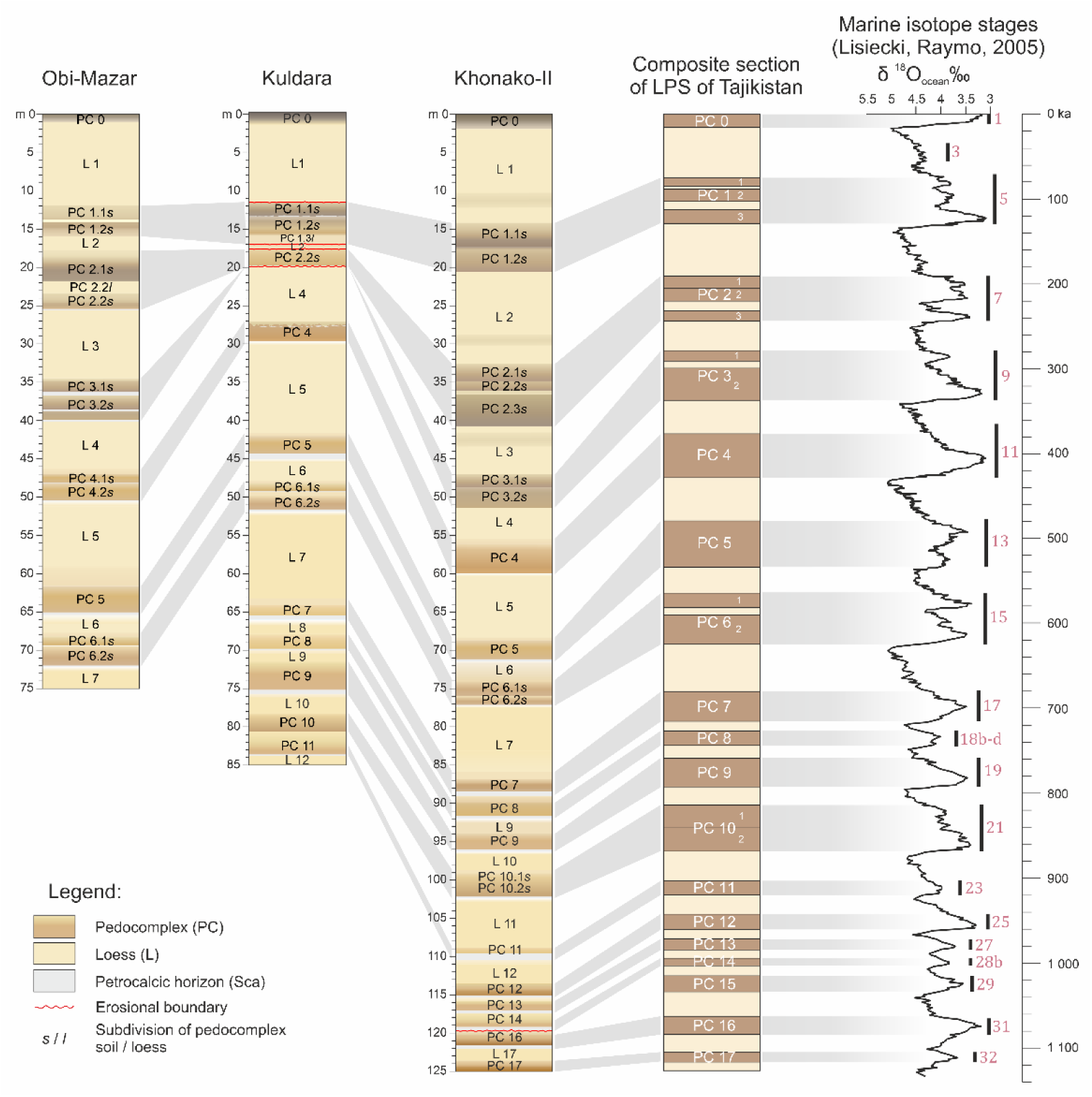
The stratigraphy of loess-paleosol sequences of OBM, KUL, and KH-II and correlations of general stratigraphy with marine isotope stages (MIS) after Dodonov and Baiguzina (1995). The δ^18^O benthic stack as a global ice volume from Lisiecki and Raymo (2005).

### 4.2 Magnetic susceptibility

The magnetic susceptibility values demonstrate higher values in paleosols than in loess units (Fig. 5). The loess horizons are quite uniform in their magnetic properties, and their mean values of magnetic susceptibility varied in the range of 2.8 to 3.0 × 10^-4^ SI units (Fig. 5), whereas in paleosols the values are, on average, 2× to 3.5× higher. Petrocalcite horizons, which are illuvial horizons of secondary precipitation of carbonates from overlying soil horizons, are characterized by low values of magnetic susceptibility due to abundant amounts of CaCO_3_ (Fig. 5). Magnetic susceptibility can be used for stratigraphic correlation of the studied sections since each PC has its characteristic features. PC 4 is characterized by one sharp peak with maximum values up to 8.4 to 10.6 × 10^-4^ SI units (Fig. 5). Sometimes, two paleosols are distinguished in the structure of the PC (for example, OBM) by the presence of a thin petrocalcic (Bkk) horizon in the middle of the PC. However, the paleosol of the final stage of formation of the pedocomplex (PC 4.1) is weakly developed with significantly lower values of magnetic susceptibility than PC 4.2, without forming a separate peak. The magnetic susceptibility of PC 5 and PC 6 is slightly lower than that of PC 4, with maximum values of 7.0 to 8.1 × 10^-4^ SI units (Fig. 5). PC 5 across the study locations was characterized by one wide peak. PC 6 consists of two paleosols, sometimes intercalated with a thin (<50 cm) loess layer affected by soil formation processes. Both paleosols of PC 6 are well distinguished by two closely spaced peaks in magnetic susceptibility, in which the values are higher for the more developed paleosol PC 6.2.

**Fig. 5.**
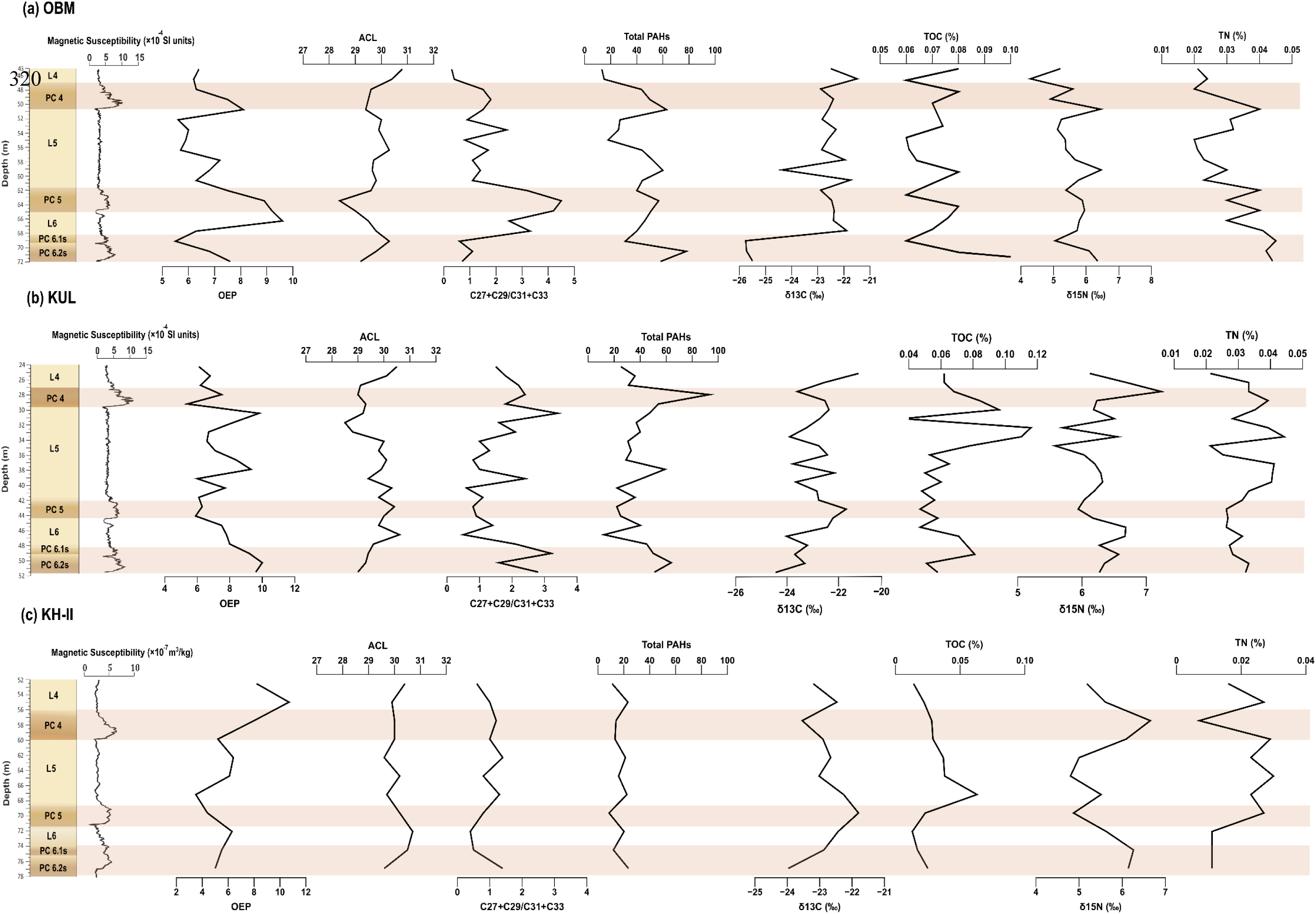
Stratigraphic profiles of magnetic susceptibility, *n*-alkane indices (OEP, ACL, and C_27_+C_29_/C_31_+C_33_ ratio), total PAH concentrations (ng/g), and stable isotopes (δ¹³C and δ¹⁵N), along with total organic carbon (TOC, %) and total nitrogen (TN, %) for the (a) OBM, (b) KUL and (c) KH-II locations. For OBM and KUL locations, field magnetic susceptibility is used (dimensionless units), for the KH-II location, laboratory measurements of mass-specific magnetic susceptibility are used (m^3^/kg). The previously published *n*-alkane proxy (ACL and C_27_+C_29_/C_31_+C_33_ ratio) and total PAHs data by Chirakkal et al. (2025a, 2025b) were used in this figure for calculation and interpretation.

### 4.3 Stable isotopes (δ¹³C and δ¹⁵N)

The δ¹³C values across all three study locations (OBM, KUL, and KH-II) range between –26‰ and –21‰, showing site-dependent patterns, with relatively enriched values typically associated with the paleosol units PC 4 and PC 5 (Fig. 5 and Table S1). At the OBM, δ¹³C values become progressively depleted from PC 5 downward to PC 6, peaking around –26‰ in PC 6, suggesting either an increased contribution from C_3_ vegetation or diagenetically altered organic matter during soil formation (Fig. 5a). However, KUL shows a similar trend, with PC 4 recording the most enriched values (–22‰), while PC 6 is isotopically lighter (–25‰) (Fig. 5b). Similarly, the KH-II mirrors this pattern, albeit with slightly less variability and a sharp peak at PC 5 (Fig. 5c). The δ¹⁵N values range from ∼4‰ to ∼7‰, showing an inconsistent trend across three profiles (Fig. 5 and Table S1). However, PC 4 shows the highest δ¹⁵N values at all three locations, reaching ∼7–8‰ at the KUL site, which could be indicative of intensified microbial processing or greater organic matter turnover under more stable soil-forming conditions. In contrast, the loess layers between the paleosols, by contrast, show lower δ¹⁵N values and more depleted δ¹³C signatures, may point to reduced biological productivity and organic input (Fig. 5). These stable isotope data of δ¹³C and δ¹⁵N reflect shifts in vegetation and nitrogen cycling.

### 4.4 *n*-Alkanes and PAHs

The GC-MS traces reveal the presence of *n*-alkanes, PAHs, and unresolved complex mixture (UCM) in the total ion chromatograms (TIC) (Fig. 6). The *n*-alkanes detected in TIC of the non-polar lipid fractions of the studied sediments comprise a homologous series ranging from C_15_ to C_33_ (Fig. 6). Please note that lipid biomarker datasets (*n*-alkanes and PAHs) have been published previously (Chirakkal et al., 2025a, 2025b) and are reused here for comparative and integrative purposes alongside newly generated stable isotope and magnetic susceptibility data. The *n*-alkane series exhibits a bimodal distribution with two distinct centers at C_16_– C_18_ and a dominant C_29_–C_31_ range, and a strong odd-over-even carbon predominance is also observed in the range of C_27_–C_33_ homologues (Fig. 6).

**Fig. 6.**
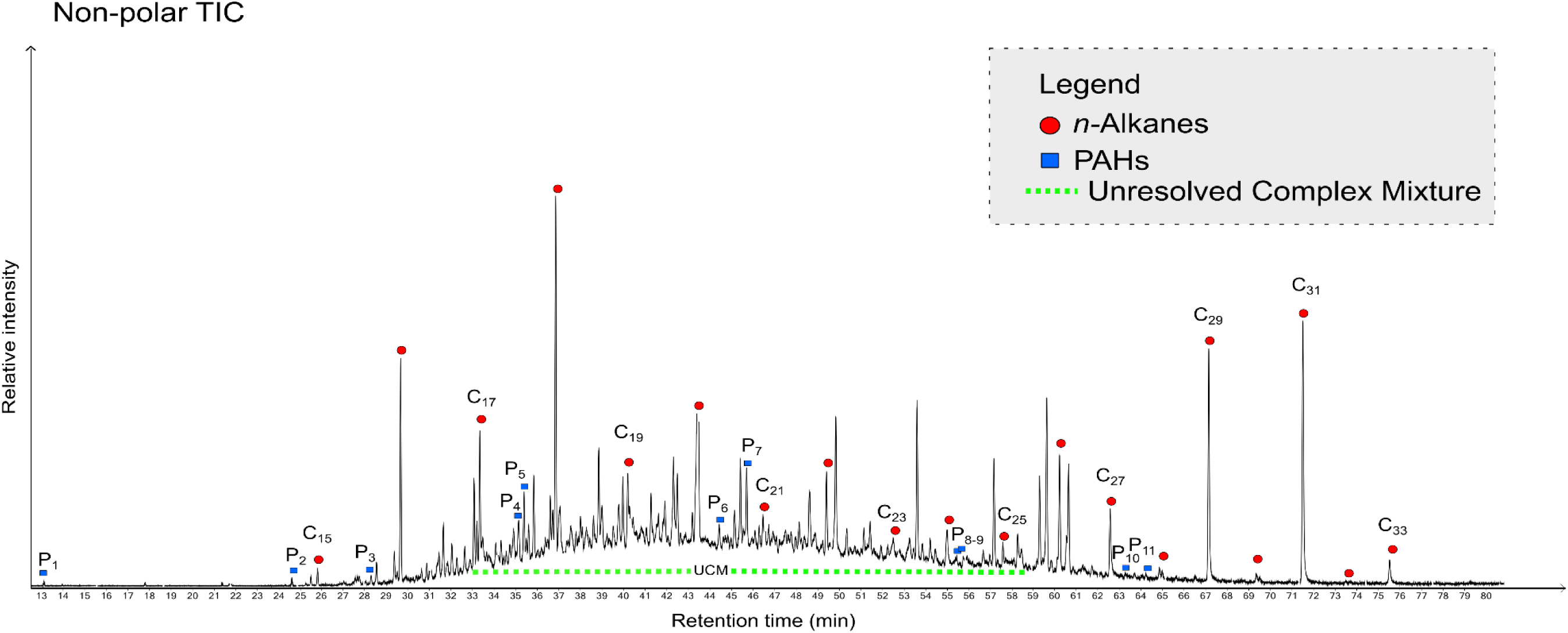
Total ion chromatogram (TIC) of the non-polar fraction of a sediment sample (ISO 45; collected from the PC 5 layer of OBM). C# shows the length of the chains of *n*-alkanes, P# shows the identified PAH compounds, and the presence of unresolved complex mixture (UCM). The *n*-alkane and PAH compound names, formulae, and spectral details can be found in Table S2.

*n*-Alkane-based indices show marked variability across stratigraphy, highlighting contrasting environmental conditions between paleosols and loess layers. The ACL values range between ∼28.5 and 30.5, with the highest values (>30) observed in loess layers and the lowest values (<29) in paleosol layers of all three locations, indicating a shift in grass and tree vegetation in glacial-interglacial cycles (Fig. 5 and Table S1). Further, the C_27_+C_29_/C_31_+C_33_ ratio exhibits a fluctuating trend with generally higher values at the OBM and KUL locations, whereas KH-II shows a relatively stable pattern with consistently lower values (Fig. 5). Similarly, the OEP index exhibits pronounced odd-over-even predominance in PC 4 and PC 5 (values >8 in OBM and KUL), consistent with well-preserved, minimally degraded, or higher plant inputs (Fig. 5 and Table S1). In contrast, at the KH-II location observed comparatively higher ACL (>29.5) and lower OEP values (∼4-6) were observed, suggesting increased grassy vegetation with lesser plant matter input in SOM (Fig. 5 and Table S1).

The PAH compounds detected in TIC of the non-polar lipid fractions of the studied sediments (Fig. 6) comprise 2-ring (e.g., Naphthalene) to 5-ring (e.g., Benzopyrene) compounds (Table S2). Total PAH concentrations are highest within PC 4 in all locations, particularly at OBM and KUL, where levels exceed 50-100 ng/g (Fig. 5a,b, and Table S1). This elevated PAH abundance in the SOM of paleosols at OBM and KUL could be the reflection of enhanced biomass burning due to anthropogenic or natural factors in the setting. On the other hand, the paleosol layers at the KH-II location display lower PAH concentrations, which may reflect a less disturbed environment (Fig. 5c and Table S1).

## 5. Discussion

### 5.1 Sources of organic matter and its preservation

The distribution and composition of *n*-alkanes across the studied profiles provide key insights into the origin and preservation state of organic matter. The diversity of *n*-alkanes found in SOM, with predominance of their long-chain homologues, and their distinctive OEP (>4) strongly suggests that they were mainly derived from terrestrial plant material (Jansen and Wiesenberg, 2017). The OEP values found in the loess-paleosol samples from the KLP sites (>5) are relatively high, though significantly lower than those reported by Zech et al. (2010) for fresh plant material (OEP = 15.0-17.9). This reduction in values is expected because OEP decreases with diagenesis and soil formation. Accordingly, our values are close to those observed for soil samples under grassland (4.3) and deciduous forest (5.0) in the same study. However, the lower OEP values observed in loess layers could be an implication of either a decreased contribution of higher-order plant material or enhanced diagenetic alteration compared to paleosols. Within the general context of basic principles of soil formation and SOM weathering (Degens, 1967; Deming and Baross, 1993), the former explanation is more likely than the latter.

A previous study by Chirakkal et al. (2025a) has highlighted the degradation and/or fire-affected organic contents in the loess and paleosol samples from the KLP locality. Also, the presence of unresolved complex mixtures (UCMs) is often interpreted as degradation of *n*-alkanes in SOM, and reported in GC-chromatograms of biodegraded petroleum or bitumen (Frysinger et al., 2003; Venkatesan and Kaplan, 1982). The δ¹³C range observed in our dataset falls within the values expected for microbially degraded sedimentary organic carbon, as reported by Wynn et al. (2005), suggesting that microbial alteration is a key source of this isotopic variability. According to Zech et al. (2007), TOC and C/N can be used as proxies for SOM degradation in loess deposits. The cross-plot analyses showed a significant positive correlation between TOC and C/N (R^2^ = 0.52) corroborates our δ¹³C interpretation (Fig. 3a). Both TOC and C/N are inversely correlated with δ^13^C (R^2^ = 0.56 and R^2^ = 0.57, respectively, see Fig. 7b and c). We conclude that most likely SOM degradation exerted a significant control on TOC, C/N and likewise on δ^13^C signatures of the KLP sequences. The observed relationships align with patterns reported by Zech et al. (2007) from the Tumara Valley, indicating a preferential loss of carbon due to decomposition in sediments over time.

**Fig. 7.**
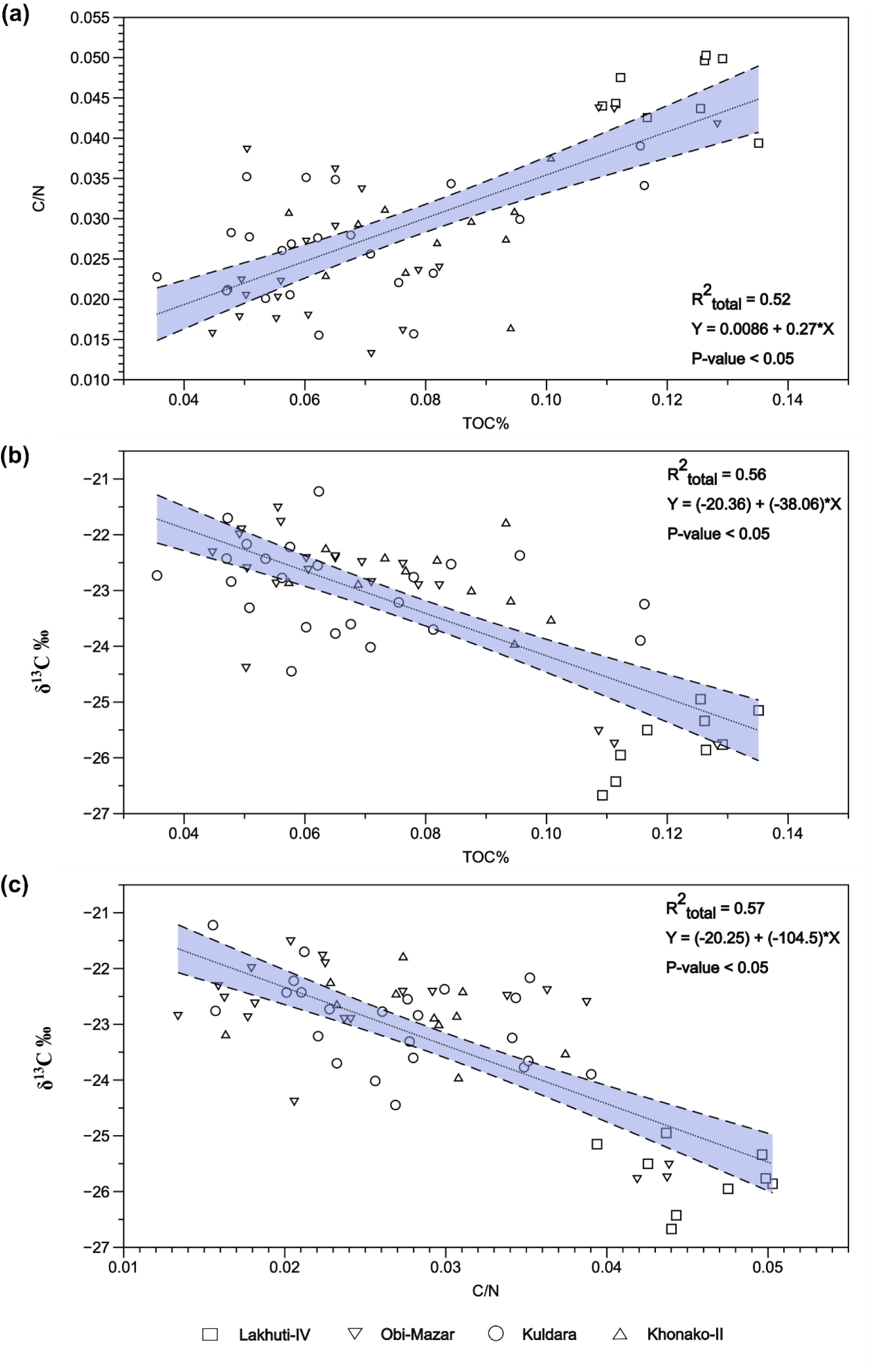
Correlation plot between δ^13^C, TOC, and C/N (n = 63): (a) δ^13^C vs. C/N (R^2^ = 0.57), (b) δ^13^C vs. TOC (R^2^ = 0.56), and (c) C/N vs. TOC (R^2^ = 0.52). The positive correlations indicate that δ^13^C in the KLP sequences is influenced by SOM degradation.

### 5.2 Paleoecological interpretations of the Paleolithic KLP

The integrated geochemical and magnetic susceptibility datasets from the KLP sequences point to dynamic vegetation and soil development patterns linked to regional climatic shifts, which may have been at least partially affected by post-burial taphonomic conditions. The δ¹³C values vary from –26 to –21‰ across the profiles, suggesting mixed vegetation inputs with an overall dominance of C₃ plants, but with localized enrichment of C_4_ (up to –21‰) at KH-II locality, hinting at increased aridity or a contribution from grass and/or herbs during its formation. In Central Asia, C_4_ vegetation grows best in warmer and drier climatic conditions and thrives with increased summer precipitation, whereas the C_3_ vegetation is more commonly associated with cooler climates and winter/spring precipitation (Yang and Ding, 2006; Zhao et al., 2022). Additionally, C_3_ grasses are common in this part of Central Asia, but due to the “canopy effect” of carbon recycling of heavier isotopes in closed vegetation environments, their isotopic signatures in open environments are similar to those of C_4_ grasses (Bonafini et al., 2013; Buchmann et al., 1997; van der Merwe and Medina, 1991). The seasonal differences in precipitation play a key role in shaping open vs. closed landscapes, with variability in moisture availability exerting a strong influence on the distribution of woodland (closed) and grassy (open) mosaics across Central Asia, as highlighted in previous studies on paleoclimate and vegetation dynamics from this region (Dodonov et al., 2006; Dodonov and Baiguzina, 1995; Yang and Ding, 2006; Zhao et al., 2022).

The broader transition towards open grassland ecosystems in the study region since 0.8 Ma is further supported by magnetic susceptibility data, where there is an observed trend of increasing magnetic susceptibility for loess-paleosol sections of Southern Tajikistan starting from PC 4 (MIS 11; Ding et al., 2002; Dodonov et al., 2006; Kulakova and Kurbanov, 2023). The synchronicity of enriched δ¹³C, high ACL values, high C_27_+C_29_/C_31_+C_33_ ratio values, and increased magnetic susceptibility in PC 4, PC 5, and PC 6 from our study supports the interpretation of a phase marked by more stable soil formation. In contrast, the loess layers separating the paleosols exhibit enriched δ¹³C, low TOC and TN contents, and depressed magnetic susceptibility, suggesting colder and more arid conditions with limited vegetation cover and soil development. The cyclical nature of these shifts between loess and paleosol formation reflects glacial–interglacial variability in Central Asia, where interglacial phases favored pedogenesis and organic accumulation, and glacial periods led to dust deposition and landscape destabilization.

Based on our *n*-alkane indices and isotope data, the KLP region hosted forest-grassland mosaics with a more tree-dominated landscape in interglacial periods and a more grass-steppe environment during the glacial periods. Moreover, the paleosols at three locations coincide with increased δ¹⁵N values, pointing to increased N losses due to accelerated mineralisation. According to Amundson et al. (2003), δ¹⁵N values tend to increase with rising mean annual temperature (MAT) and decrease with higher mean annual precipitation (MAP). Applying this model to the KLP sediments suggests that phases of soil development were associated with an open nitrogen cycle and δ¹⁵N enrichment, likely driven by more robust evaporitic conditions, such as warmer and drier climates (Figueroa and Davy, 1991; Porqueddu et al., 2016). A modern analogue may be found in Mediterranean climates, where cool, wet winters and hot, dry summers limit tree growth but still support productive grasslands. This climatic seasonality with moisture concentrated in the cooler months may explain the co-existence of inhibited arboreal expansion and active pedogenesis during these phases.

The paleoclimatic trends documented in regional and global proxy records further support our interpretation, indicating enhanced soil development and vegetation expansion during Marine Isotope Stages (MIS) 11, 13, and 15. These interglacial intervals coincide with peaks in speleothem δ¹⁸O values from East Asia (Cheng et al., 2016; Liu et al., 2020) and with reduced dust fluxes in the Chinese Loess Plateau (Guo et al., 2009; Qiang et al., 2011), reflecting warmer and more humid conditions conducive to pedogenesis. Such climatic phases likely correspond to the development of the paleosols PC 4, PC 5, and PC 6 in the KLP sequence. The onset of environmental transitions inferred from our geochemical proxies (δ¹³C, δ¹⁵N, TOC, TN, and magnetic susceptibility) aligns with these broader interglacial trends, suggesting that the KLP region has experienced favorable ecological conditions that may have facilitated hominin presence or occupation during these periods, as also indicated by the higher density of archaeological materials reported from the KLP region of Southern Tajikistan (Anoikin et al., 2023; Ranov, 1995; Schäfer et al., 1998). While the possibility exists of diagenetically altered SOM attenuating climate signals expressed in the bulk stable isotope data, previous studies from the same region (Chirakkal et al., 2025a, 2025b; Dodonov et al., 2006; Dodonov and Baiguzina, 1995; Häggi et al., 2019; Mestdagh et al., 1999; Yang and Ding, 2006) consistently indicate that soil formation and forest development were strongest during interglacial periods and weakest during glacial periods, supporting the overall interpretation presented above.

### 5.3 Human-environment interaction during the Early Paleolithic

The integration of lithic artifact counts and PAH concentrations reveals a distinct spatial gradient of human-environment interaction across the KLP landscape. OBM is located directly on the bank of the Obi-Mazar River. The exceptionally high number of archaeological materials (over 2,000 in PC 4 and several hundred in PC 5 and PC 6) indicates a frequent on-site occupation area (Fig. 8). These numbers coincide with the highest total PAH concentrations (48.1 ng/g in PC 4 and 52.8 ng/g in PC 5), suggesting frequent burning events or continuous fire use (Fig. 8). At the same stratigraphic levels, lithic concentrations are also highest, and the paleosol horizons display enhanced pedogenic development with higher magnetic susceptibility values highlighting of stable surface conditions conducive to repeated occupation. The co-occurrence of these proxies with elevated PAHs abundance, indicating biomass combustion, dense lithic assemblages denoting repeated tool production or use, and stable paleosol formation reflecting extended landscape stability, which supports the interpretation that OBM served as a recurrent activity area where hominins repeatedly exploited the riverine environment during favorable climatic intervals. The continuity of more closed canopy vegetation around the Obi-Mazar River may signal hominin preference for retaining sheltered occupation zones amongst tracts of grassier areas where foraging opportunities would have been more abundant. Furthermore, the potential repeated presence of hominins in the surrounding landscape may have influenced local faunal behavior, possibly leading grazing animals to avoid frequently hominin used areas, a pattern conceptually comparable to the “Ecology of Fear” (*sensu* Brown et al., 1999). Although this remains a tentative interpretation due to the limited archaeological sample size, it provides a plausible hypothesis that can be further evaluated through future zooarchaeological and spatial analyses.

**Fig. 8.**
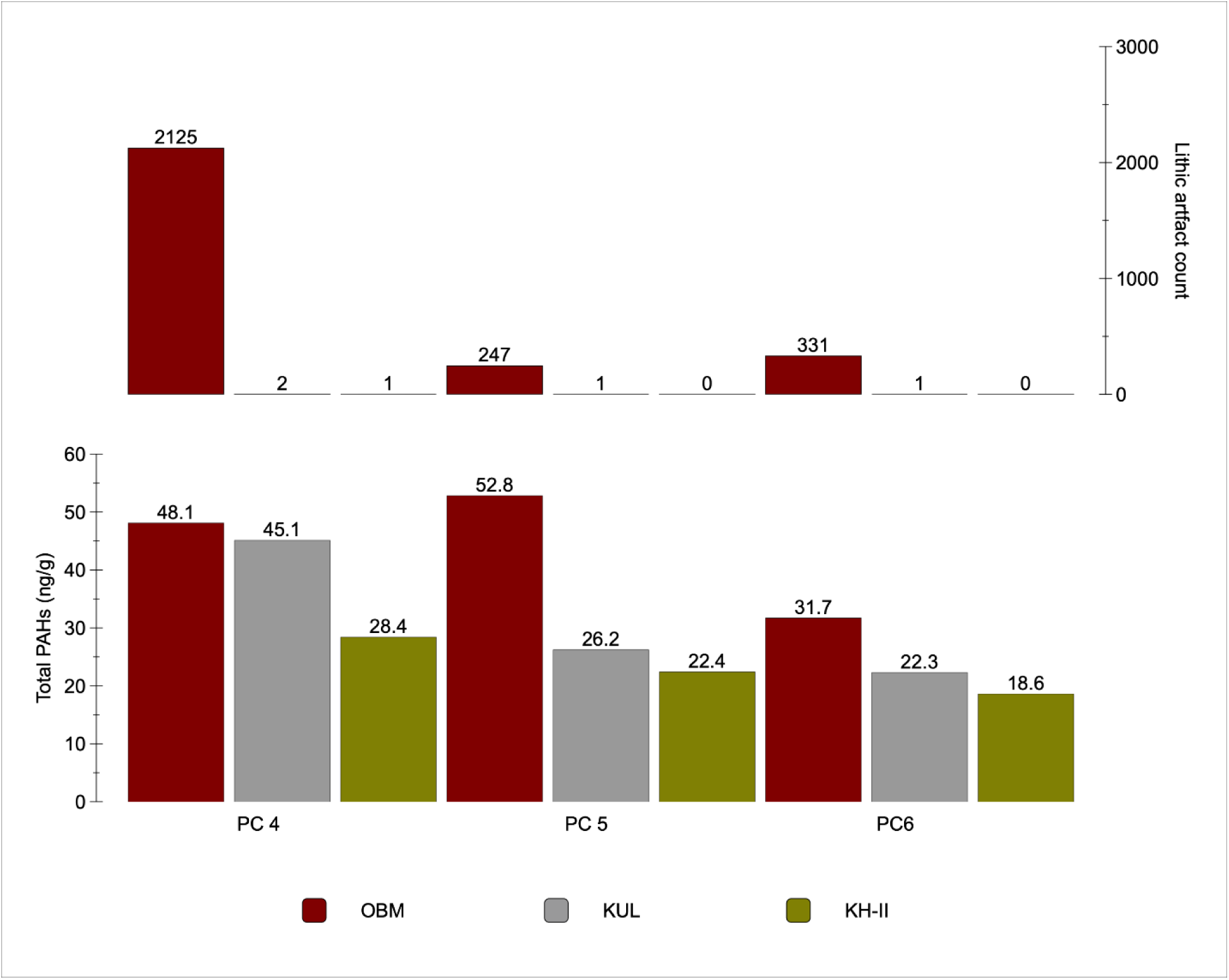
Lithic artifact counts and total PAH concentrations for PC 4, 5, and 6 at OBM, KUL, and KH-II study location of the KLP. The previously published PAHs data by Chirakkal et al. (2025a, 2025b) were used in this figure for interpretation.

In contrast, KUL, situated about 2 km from the Obi-Mazar, shows moderate PAH levels (45.1 ng/g in PC 4, decreasing through PC 5 and PC 6) but only minimal lithic artifacts, suggesting a near-site setting with evidence of regional fire signals and more limited occupation or human activity (Fig. 8). Moreover, KH-II, which is located approximately 15 km from the river, has the lowest PAH concentrations (28.4 ng/g in PC 4 decreasing to 18.6 ng/g in PC 6) and negligible lithic material, representing an off-site background signal with little to no direct human influence in the immediate area (Fig. 8). Therefore, the KH-II location might have remained as natural landscapes with more C_4_ or grassy C_3_ vegetation and minimal human impact, which is reflected in these geochemical fingerprints that suggest lower fire biomarker and geochemical input in SOM in tune with climate, rather than hominin-driven selection pressures. Together, these data indicate that hominin influence in the form of fire- and vegetation pressure was most pronounced along the riverbank and progressively diminished with increasing distance from it. At the riverine locality (OBM), elevated PAH concentrations, higher TOC values, and abundant lithic materials point to recurrent burning and occupation within stable, well-developed paleosols that provided access to water, raw materials, and vegetation. In contrast, the more distal site (KH-II) shows markedly lower PAH concentrations and negligible archaeological evidence, suggesting limited or no human activity. This spatial gradient supports a model in which hominin presence was concentrated in ecologically favorable riverine settings and declined toward the upland loess surfaces, where environmental resources and soil stability were comparatively restricted.

## 6. Conclusion

This study integrates biomarker indices, magnetic susceptibility, stable isotope geochemistry, and archaeological evidence from loess-paleosol sequences in the KLP of southern Tajikistan to reconstruct paleoenvironmental dynamics and human-environment interactions during the Early Paleolithic. The *n*-alkane distributions with high OEP and ACL values indicate that the sedimentary organic matter is primarily derived from higher-order terrestrial plants, with better preservation within paleosols than in loess layers. The cross-plots between δ¹³C, TOC, and C/N further reveal that part of the isotopic variability is controlled by SOM degradation, consistent with patterns observed in other loess regions. Elevated δ¹⁵N values in paleosols suggest phases of increased mineralization under warmer and drier conditions, while magnetic susceptibility trends and δ¹³C values document a long-term shift toward more open grassland ecosystems since ∼0.8 Ma. Moreover, the magnetic susceptibility, *n*-alkane indices, and δ¹³C records from PCs 4, 5, and 6 appear to broadly correspond with interglacial phases (MIS 11, 13, and 15) identified in regional paleoclimate archives, which were characterized by warmer and more humid conditions that likely promoted the expansion of mixed forest-grassland vegetation.

Superimposed on these environmental trends are spatial variations that likely reflect differences in the intensity of hominin presence across the landscape. At the riverine locality (OBM), relatively high lithic artifact densities and elevated PAH concentrations suggest repeated or sustained activity, possibly involving fire use within a resource-rich setting. In contrast, the near-site area (KUL) shows moderate PAH abundances but limited lithic evidence, implying more transient or episodic use, whereas the off-site locality (KH-II) records low PAH values and negligible archaeological material, indicating minimal or no detectable human impact. Collectively, these patterns suggest that hominin activity was more frequent in ecologically favorable localities near riverine corridors. The multi-proxy approach adopted here not only refines the paleoecological context of the KLP region but also provides a framework for understanding how early hominins responded to and modified their environments in Central Asia during the Pleistocene.

## CRediT authorship contribution statement

**Aljasil Chirakkal:** Writing – original draft, Writing – review & editing, Conceptualization, Methodology, Validation, Formal analysis, Data curation, Investigation, Visualization. **Ekaterina Kulakova:** Writing – review & editing, Investigation, Data curation, Formal analysis. **Jago J. Birk:** Writing – review & editing, Supervision. **Calin C. Steindal:** Writing – review & editing, Supervision. **Anton Anoikin**: Writing – review & editing, Data curation, Investigation. **Redzhep Kurbanov**: Data curation, Investigation. **Piotr Sosin**: Data curation, Investigation. **Peixian Shu**: Data curation, Investigation. **David K. Wright:** Writing – review & editing, Conceptualization, Methodology, Formal analysis, Validation, Supervision, Project administration, Funding acquisition, Resources.

## Acknowledgments

This research is supported by the Nordforsk (Project number: 105204) grant awarded to Dr. Jan-Pieter Buylaert (PI of the THOCA project, Technical University of Denmark, DTU). We thank Olga Meshcheryakova and Olga Tokareva for conducting field magnetic susceptibility measurements. We gratefully acknowledge the helpful support of Ankit Yadav for his valuable suggestion on the lipid biomarker discussion. We want to thank Nadya ‘Lennie’ Wright, who helped us to prepare isotope samples, and William Hagopian (University of Oslo, Norway) and Pål Tore Mørkved (University of Bergen, Norway) for analyzing our samples in their laboratory. We are grateful to Redzhep Kubanov and Anton Anoikin as well as all of the contributing members of the field crews in Tajikistan for leading the archaeological investigations upon which our microarchaeological approach is built.

