## Supplemental for "Geochemical records from loess sediments provide insights into early hominin influence on the landscape in the Khovaling region of Southern Tajikistan, Central Asia"

**Supplemental Figure S1:**

(a) Average annual temperature (°C) and (b) average annual precipitation (mm) data of Khovaling region of Tajikistan (source: Temperature and Precipitation information (2003-2023), https://crudata.uea.ac.uk/cru/data/hrg/). The red dot denotes the location of the KLP.

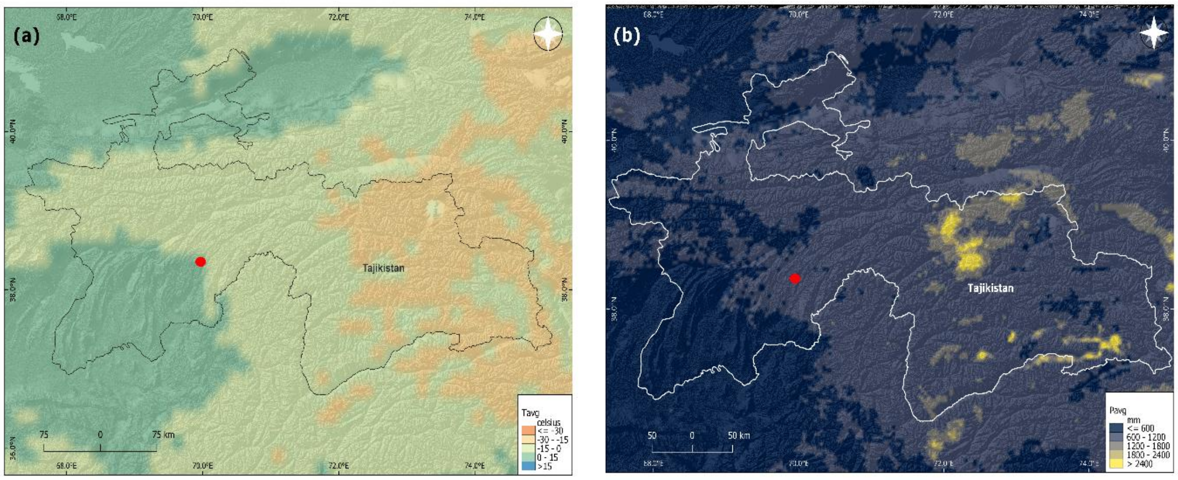


**Supplemental Text S1:**

The stable carbon ($\delta^{13}C$) and nitrogen ($\delta^{15}N$) isotopes samples were analyzed at the CLimate Interpretation of Plant Tissue (CLIPT) laboratory in the Department of Biosciences at the University of Oslo. Due to the financial crisis, CLIPT Laboratory was shut down in 2023. Therefore, we moved laboratory analyses to the Facility for advanced isotopic research and monitoring of weather, climate, and biogeochemical cycling (FARLAB) at the University of Bergen. However, both laboratories followed identical protocols to keep consistent measurements.

### Supplemental Table S1:

The results of OEP, ACL, $C_{27}$+$C_{29}$/$C_{31}$+$C_{33}$, total PAHs (ng/g) TOC (%), TN (%), $\delta^{13}C_{PDB}$ (‰), and $\delta^{15}N_{AIR}$ (‰) of samples analyzed in this study. The previously published *n*-alkane and PAHs data by Chirakkal et al. (2025a, 2025b) were used in this figure for interpretation.

| Sample ID | Study site | Pedocomplex | Depth (m) | OEP | ACL | $C_{27}$+$C_{29}$/$C_{31}$+$C_{33}$ | Total PAHs (ng/g) | $\delta^{13}C_{PDB}$ (‰) | TOC (%) | $\delta^{15}N_{AIR}$ (‰) | TN (%) |
|---|---|---|---|---|---|---|---|---|---|---|---|
| ISO 59 | OBM | L4 | 45 | 6.4 | 30.8 | 0.3 | 13.2 | -22.5 | 0.08 | 5.2 | 0.02 |
| ISO 57 | OBM | L4 | 46 | 6.2 | 30.4 | 0.4 | 15.0 | -21.5 | 0.06 | 4.3 | 0.02 |
| ISO 56 | OBM | PC 4 | 47.5 | 6.3 | 29.6 | 1.5 | 43.6 | -22.9 | 0.08 | 5.6 | 0.02 |
| ISO 55 | OBM | PC 4 | 48.2 | 7.5 | 29.5 | 1.8 | 50.0 | -22.4 | 0.07 | 4.9 | 0.03 |
| ISO 54 | OBM | PC 4 | 49.8 | 8.1 | 29.4 | 1.5 | 62.8 | -22.6 | 0.07 | 6.4 | 0.04 |
| ISO 53 | OBM | L5 | 51 | 5.6 | 30.0 | 0.9 | 27.1 | -22.8 | 0.07 | 5.2 | 0.03 |
| ISO 52 | OBM | L5 | 53 | 6.0 | 29.9 | 2.4 | 26.0 | -22.3 | 0.06 | 5.1 | 0.03 |
| ISO 51 | OBM | L5 | 56 | 5.9 | 30.1 | 0.8 | 18.0 | -22.6 | 0.06 | 5.4 | 0.02 |
| ISO 50 | OBM | L5 | 59 | 5.7 | 30.3 | 1.7 | 44.0 | -22.9 | 0.06 | 5.4 | 0.02 |
| ISO 49 | OBM | PC 5 | 61.8 | 7.2 | 29.7 | 1.1 | 52.3 | -22.0 | 0.08 | 5.7 | 0.02 |
| ISO 42 | OBM | PC 5 | 62 | 6.8 | 29.7 | 1.4 | 60.0 | -24.4 | 0.07 | 6.5 | 0.03 |
| ISO 43 | OBM | PC 5 | 62.3 | 6.3 | 29.8 | 1.1 | 44.3 | -21.7 | 0.06 | 5.8 | 0.02 |
| ISO 44 | OBM | PC 5 | 62.6 | 7.5 | 29.6 | 3.2 | 40.0 | -22.9 | 0.08 | 5.4 | 0.04 |
| ISO 45 | OBM | PC 5 | 62.8 | 8.9 | 28.4 | 4.5 | 56.8 | -22.5 | 0.07 | 5.9 | 0.03 |
| ISO 46 | OBM | PC 5 | 63 | 9.2 | 29.0 | 4.2 | 51.0 | -22.4 | 0.07 | 5.9 | 0.04 |
| ISO 47 | OBM | PC 5 | 63.6 | 9.6 | 29.5 | 2.5 | 46.0 | -22.4 | 0.06 | 5.8 | 0.03 |
| ISO 48 | OBM | PC 5 | 64 | 6.3 | 29.8 | 3.3 | 40.0 | -21.9 | 0.08 | 5.7 | 0.04 |
| ISO 121 | OBM | L6 | 66 | 5.5 | 30.3 | 0.6 | 31.1 | -25.8 | 0.13 | 5.1 | 0.04 |
| ISO 122 | OBM | PC 6 | 69 | 6.8 | 29.8 | 1.1 | 77.9 | -25.7 | 0.11 | 6.1 | 0.04 |
| ISO 123 | OBM | PC 6 | 71 | 7.6 | 29.2 | 0.7 | 58.2 | -25.5 | 0.11 | 6.3 | 0.04 |
| ISO 27 | KUL | L4 | 24.2 | 6.1 | 30.5 | 1.5 | 25.1 | -21.2 | 0.06 | 5.6 | 0.02 |
| ISO 7 | KUL | PC 4 | 27.3 | 6.8 | 30.1 | 1.8 | 36.0 | -22.6 | 0.06 | 6.2 | 0.03 |
| ISO 8 | KUL | PC 4 | 27.5 | 6.2 | 29.1 | 2.2 | 31.0 | -23.6 | 0.07 | 6.7 | 0.03 |
| ISO 9 | KUL | PC 4 | 27.8 | 7.5 | 29.0 | 2.4 | 93.1 | -22.5 | 0.08 | 5.7 | 0.03 |
| ISO 10 | KUL | PC 4 | 28 | 5.4 | 29.3 | 1.8 | 54.0 | -22.4 | 0.10 | 5.7 | 0.03 |
| ISO 11 | KUL | PC 4 | 28.3 | 9.8 | 29.2 | 3.4 | 47.0 | -22.7 | 0.04 | 6.0 | 0.02 |
| ISO 12 | KUL | PC 4 | 28.6 | 8.2 | 28.5 | 1.6 | 37.0 | -23.2 | 0.12 | 5.2 | 0.03 |
| ISO 13 | KUL | PC 4 | 29.2 | 6.7 | 28.8 | 2.1 | 40.0 | -23.9 | 0.12 | 6.1 | 0.04 |
| ISO 14 | KUL | L5 | 36 | 6.6 | 30.0 | 1.0 | 30.5 | -22.8 | 0.08 | 5.1 | 0.02 |
| ISO 1 | KUL | PC 5 | 42.1 | 7.1 | 29.8 | 1.3 | 33.0 | -22.4 | 0.05 | 5.5 | 0.02 |
| ISO 2 | KUL | PC 5 | 42.3 | 8.4 | 30.1 | 0.8 | 29.0 | -23.8 | 0.07 | 5.7 | 0.03 |
| ISO 3 | KUL | PC 5 | 42.5 | 9.3 | 29.9 | 1.0 | 59.0 | -22.2 | 0.05 | 5.8 | 0.04 |
| ISO 4 | KUL | PC 5 | 43 | 6.0 | 29.4 | 2.4 | 41.0 | -23.7 | 0.06 | 5.8 | 0.04 |
| ISO 5 | KUL | PC 5 | 43.5 | 7.7 | 30.3 | 0.6 | 22.2 | -22.8 | 0.05 | 5.7 | 0.03 |
| ISO 6 | KUL | PC 5 | 44 | 6.1 | 29.8 | 1.1 | 36.0 | -22.8 | 0.06 | 5.5 | 0.03 |
| ISO 19 | KUL | L6 | 46 | 6.3 | 30.4 | 0.8 | 22.0 | -21.7 | 0.05 | 5.4 | 0.02 |
| ISO 20 | KUL | L6 | 47 | 5.9 | 30.0 | 0.9 | 24.9 | -22.2 | 0.06 | 5.7 | 0.02 |
| ISO 21 | KUL | PC 6 | 48.4 | 7.5 | 29.8 | 1.4 | 40.0 | -22.4 | 0.05 | 6.2 | 0.02 |
| ISO 22 | KUL | PC 6 | 49 | 7.8 | 30.6 | 0.5 | 12.1 | -24.0 | 0.07 | 6.2 | 0.03 |
| ISO 23 | KUL | PC 6 | 49.5 | 8.0 | 29.6 | 2.1 | 45.0 | -23.2 | 0.08 | 5.8 | 0.02 |
| ISO 24 | KUL | PC 6 | 50 | 9.2 | 29.4 | 3.2 | 50.0 | -23.7 | 0.08 | 6.1 | 0.02 |
| ISO 25 | KUL | PC 6 | 50.6 | 10.0 | 29.3 | 1.6 | 63.8 | -23.3 | 0.05 | 5.9 | 0.03 |
| ISO 26 | KUL | PC 6 | 51 | 9.6 | 29.0 | 2.8 | 51.0 | -24.4 | 0.06 | 5.8 | 0.03 |
| ISO84 | KH-II | L4 | 54 | 8.2 | 30.4 | 0.6 | 11.0 | -23.2 | 0.01 | 5.2 | 0.02 |
| ISO78 | KH-II | PC 4 | 56.5 | 10.7 | 29.9 | 1.0 | 23.2 | -22.5 | 0.02 | 5.6 | 0.03 |
| ISO77 | KH-II | PC 4 | 58 | 8.0 | 30.0 | 1.2 | 14.0 | -23.5 | 0.03 | 6.7 | 0.01 |
| ISO76 | KH-II | PC 4 | 59 | 5.2 | 30.0 | 1.0 | 13.2 | -22.9 | 0.03 | 6.1 | 0.03 |
| ISO75 | KH-II | L5 | 62 | 6.4 | 29.6 | 1.4 | 21.3 | -22.7 | 0.04 | 5.0 | 0.02 |
| ISO74 | KH-II | L5 | 66 | 6.1 | 30.2 | 0.8 | 16.0 | -23.0 | 0.04 | 4.8 | 0.03 |
| ISO73 | KH-II | PC 5 | 70 | 3.5 | 29.7 | 1.3 | 22.4 | -22.3 | 0.06 | 5.5 | 0.02 |
| ISO72 | KH-II | L6 | 72.6 | 4.4 | 30.2 | 0.8 | 8.6 | -21.8 | 0.02 | 4.9 | 0.03 |
| ISO71 | KH-II | PC 6 | 75 | 6.3 | 30.7 | 0.4 | 20.2 | -22.4 | 0.01 | 5.6 | 0.01 |
| ISO70 | KH-II | PC 6 | 76 | 5.5 | 30.5 | 0.5 | 12.0 | -22.9 | 0.02 | 6.3 | 0.01 |
| ISO69 | KH-II | PC 6 | 77 | 5.0 | 29.6 | 1.4 | 23.6 | -24.0 | 0.02 | 6.1 | 0.01 |

**Supplemental Table S2:**

List of *n*-alkanes and polycyclic aromatic hydrocarbons (PAHs) and corresponding diagnostic ions. Where "n" equals to carbon chain length, which ranges from 15 to 33 for *n*-alkanes.

| Serial No. | Compound name | Formula | No of rings | Diagnostic ions (m/z) |
|---|---|---|---|---|
| 1 | *n*-alkanes | $C_nH_{2n+2}$ | - | 57 |
| 2 | Naphthalene (Nap) | $C_{10}H_8$ | 2 | 128 |
| 3 | Acenaphthene (Ace) | $C_{12}H_{10}$ | 3 | 154 |
| 4 | Acenaphthylene (Acy) | $C_{12}H_8$ | 3 | 152 |
| 5 | Fluorene (Fle) | $C_{13}H_{10}$ | 3 | 166 |
| 6 | Phenanthrene (Phe) | $C_{14}H_{10}$ | 3 | 178 |
| 7 | Anthracene (Ant) | $C_{14}H_{10}$ | 3 | 178 |
| 8 | Fluoranthene (Fla) | $C_{16}H_{10}$ | 4 | 202 |
| 9 | Pyrene (Py) | $C_{16}H_{10}$ | 4 | 202 |
| 10 | Benz[a]anthracene (BaA) | $C_{18}H_{12}$ | 4 | 228 |
| 11 | Chrysene (Chr) | $C_{18}H_{12}$ | 4 | 228 |
| 12 | Benzo[b]fluoranthene (BbF) | $C_{20}H_{12}$ | 5 | 252 |
| 13 | Benzo[a]pyrene (BaP) | $C_{20}H_{12}$ | 5 | 252 |
